# Single-cell variations of circadian clock and immune gene expression in microglia and neurodegeneration

**DOI:** 10.1101/2024.06.16.599193

**Authors:** Koliane Ouk, Müge Yalçin, Ludovica Rigat, Hadas Keren-Shaul, Eyal David, Anniki Knop, Chotima Böttcher, Achim Kramer, Ido Amit, Angela Relógio, Josef Priller

## Abstract

We investigated the diurnal rhythmicity of gene expression in microglia, the resident macrophages of the brain, in health and disease. Using RNA sequencing and single-cell analysis by RNAscope, we examined wild-type mice and the R6/2 transgenic mouse model of Huntington’s disease (HD). Our findings suggest context-dependent rhythmic gene expression in microglia, exhibiting substantial variability between individual cells and brain regions over 24 hours. Notably, we observed loss of rhythmic gene expression of key clock genes in microglia from symptomatic but not presymptomatic R6/2 mice. Moreover, we identified *de novo* 24-hour rhythmic gene expression and altered diurnal patterns of immune-related genes associated with neurodegenerative diseases in microglia from symptomatic R6/2 mice. Our findings suggest circadian reprogramming of microglia in the context of neurodegeneration.

## Introduction

Microglia are the tissue-resident macrophages of the brain, first described in 1919 by del Río-Hortega, and they constitute ∼5-20% of the neuroglia. Microglia invade the CNS early in development (embryonic day 9.5 in mice) and they originate from a pool of erythromyeloid progenitors in the yolk sac^1^. Microglia are the main immune defence of the CNS and they continuously screen the microenvironment for pathological events. When a change in brain homeostasis occurs, they transition from the surveilling state to a variety of activated states. The activation process is highly context-dependent and induces a heterogeneity of microglial phenotypes. Microglia can then influence the CNS response to the stressful agent or pathological event by the release of diverse substances, including cytokines, chemokines, and growth factors. They also have a phagocytic function and help orchestrate the immunological response by interacting with infiltrating immune cells^2^. Thus, microglia play a critical role in brain development, physiological function, and disease^3, 4^.

Circadian rhythms are generated by the mammalian circadian clock, comprising a hierarchy of oscillators with the central clock located in the suprachiasmatic nucleus of the hypothalamus. The circadian clock is an endogenous and autonomous time-keeping mechanism that generates 24-hour rhythms in gene and protein expression, and thereby modulates and adjusts physiological responses and behaviour to the external geophysical time^5^. At the cellular level, circadian rhythms are generated by transcription-translation feedback loops involving the transcriptional activators *Clock, Npas*, *Arntl* (also known as *Bmal1*), *Rora/b/c,* and the transcriptional repressors *Period* (*Per1/2/3), Cryptochrome (Cry1/2/3) and Nr1d1/2* (also known as *Rev-erb-*α*/ Rev-erb-*β) and corresponding proteins. Remarkably, NPAS2 can functionally replace CLOCK in the circadian clock of mice^6^.These positive and negative elements compose the core clock network. They are expressed in most cells and confer a roughly 24-hour oscillation in the expression of so-called “clock-controlled genes” that are involved in a variety of core clock output pathways and influence physiological processes, including immune responses, metabolism, neurotransmitter release, phagocytosis and clearance of debris^7, 8, 9^.

The last decade has seen major advances in our understanding of the clock control of immunity, with strong evidence that immune responses are under the influence of the circadian clock^10–13^. Recent studies have demonstrated that glial cells also express oscillating functions critical to maintain overt circadian rhythms throughout the body^14, 15^. The presence of clock genes in microglia was first demonstrated in microglia isolated from the neonatal mouse brain and in the BV-2 microglia cell line^16^. The existence of an intrinsic molecular clock in microglia was later demonstrated by diurnal fluctuations in *Per1*, *Per2* and *Nr1d1* expression^17^. The expression of microglial *Arntl*, which is necessary to ensure circadian behaviours, has been shown to be involved in cognitive and metabolic challenges^18^. Furthermore, the time of day greatly influences the magnitude of immune responses by microglia following brain injury or bacterial challenge^19^, suggesting that endogenous circadian rhythms in microglia modulate their immune functions.

The immune and circadian systems are interconnected bidirectionally^20^. For example, the microglial clock regulates the inflammatory response to bacterial lipopolysaccharide via an *Arntl*-dependent increase in interleukin (IL)-6 expression^21^, in correlation with a greater magnitude of pro-inflammatory cytokine production^22^. *Rev-erb-*α also influences innate immunity in macrophages through the repression of *Ccl2* expression^23^. Conversely, pro-inflammatory cytokines released by microglia such as IL-1 and tumor necrosis factor (TNF) play an important role in the regulation of sleep^24^ and are able to phase-shift activity rhythms when administered at the beginning of the resting phase^25^. Disruptions of the circadian clock and immune systems are both important features in neurodegenerative diseases, such as Huntington’s disease (HD), an autosomal dominantly inherited neurodegenerative disease that results in motor, cognitive and psychiatric disturbances^26^. HD is caused by a single mutation on chromosome 4, leading to a CAG trinucleotide repeat expansion in the *Huntingtin* (*Htt*) gene. The CAG repeat encodes the amino acid glutamine, and the expansion of this repeat leads to the production of a toxic mutant protein (mHTT) with an expanded polyglutamine tract^26^. There is evidence of early immune system activation and altered immune responses in HD, with reports of activated microglia, reactive astrocytes, and changes in circulating cytokine levels^27^. Circadian behavioural abnormalities in people with HD have first been identified using actigraphy, which revealed disrupted rest-activity profiles with increased night/day activity ratios^28^. This diurnal activity disruption is replicated in the R6/2 mouse model of HD which expresses a transgene that contains exon 1 of the human *Huntingtin* gene with a large expanded CAG repeat^28^. The model is known for its early onset and rapid progression of motor and behavioural deficits. The reported daily disruption correlates with a dysregulated clock gene expression of *Per2* and *Arntl* in SCN and striatum^28^. Abnormal circadian rhythms have also been reported in mouse models of HD that expresses the full length *Htt* gene with mutant exon1, such as the Q175 mice^29^ and the BACHD mice^30^, the latter also showing dampened heart rate and body temperature rhythms^30^. Neurodegeneration with progressive atrophy of striatum and cortex, as well as microglial and astroglial activation are neuropathological hallmarks of HD^31^. HD pathology spreads progressively to other brain areas such as hypothalamus, thalamus and brain stem^32^. Accumulated evidence suggests that rather than being only the consequence of neurodegenerative and inflammatory processes, disrupted circadian clocks may directly influence key pathways involved in these two events. Since depleting *mHtt* selectively from astrocytes rescued HD neuropathology and behavioural performance^33^, emerging research focused on investigating the potential of glial cells as targets for therapeutic intervention in HD. The role of microglia in HD has not been fully explored, but there is evidence for a contribution of microglia to the pathogenesis of HD^34,35, 36^.

In a mouse model of Alzheimer’s disease (AD), an impaired microglial clock is involved in the induction of chronic neuroinflammatory alterations^37^. Moreover, disease-associated microglia states have been identified in AD transgenic mice^38,39^. Manipulations of microglia were shown to affect disease progression^4^: for example, recent studies have shown that deletion of transmembrane immune signalling adaptor (TYROBP), a microglia-enriched protein, is beneficial in both AD and HD models by reducing pro-inflammatory pathways.

To address the role of circadian rhythmicity of microglia in health and disease, we performed a circadian analysis at the transcriptomic level using bulk RNA sequencing (RNA-seq) of healthy microglia collected over a 24-hour time course. We then analyzed the impact of disease on the microglial diurnal rhythms by comparing with microglia from wild-type (WT) mice with microglia from R6/2 transgenic mice at two different stages of disease: presymptomatic (6 weeks) and symptomatic (15 weeks). We identified, in symptomatic R6/2 mice, a disrupted 24-hour rhythmicity in the expression of microglial core clock genes including *Arntl*, *Cry2*, *Nr1d2* and *Per2* but also the emergence of 24-hour rhythmicity in immune-relevant genes, such as *Interferon regulatory factor 7* (*Irf7*), a differentially expressed gene in an *in vitro* model of HD^40^, or *Intercellular adhesion molecule 2* (*Icam2)*, whose proteins are ligands for the leukocyte adhesion protein LFA-1 that mediates cellular interactions important for immune responses. We further investigated microglia at the single-cell level in cortex and striatum, the most affected structures in HD^41^, by analyzing the differential expression of the core clock gene *Arntl* and immuno-relevant genes *Ccl2* and *Irf7* using RNAscope. Our single-cell analysis confirmed in R6/2 mice a significant loss of rhythmicity for *Arntl* in cortex versus striatum. Moreover, we observed that circadian rhythms vary between individual microglia and across brain regions in WT mice.

## Results

### Circadian dysregulation in microglia from R6/2 transgenic mice

In this study, we characterized diurnal rhythms in microglia from R6/2 mice and WT littermates collected from whole brain every 4 hours across 24 hours (ZT2, 6, 10, 14, 18, 22 and 26) using bulk RNA-seq (**Figure 1A**; **Figure 1B** shows the RNAscope workflow). In the 6-week-old group, the rhythmicity analysis identified 1901 genes as significantly 24-hour rhythmic in WT mice, and 1314 genes as significantly 24-hour rhythmic in age-matched R6/2 mice (**Figures 1C-D**). Phase distribution of 24-hour oscillating genes in WT mice showed a bimodal distribution (**Figure 1C, middle and right panel**), corresponding to dawn and dusk, whereas in R6/2 mice, the phases showed a multimodal distribution (**Figure 1D, middle and right panel**).

**Figure 1.**
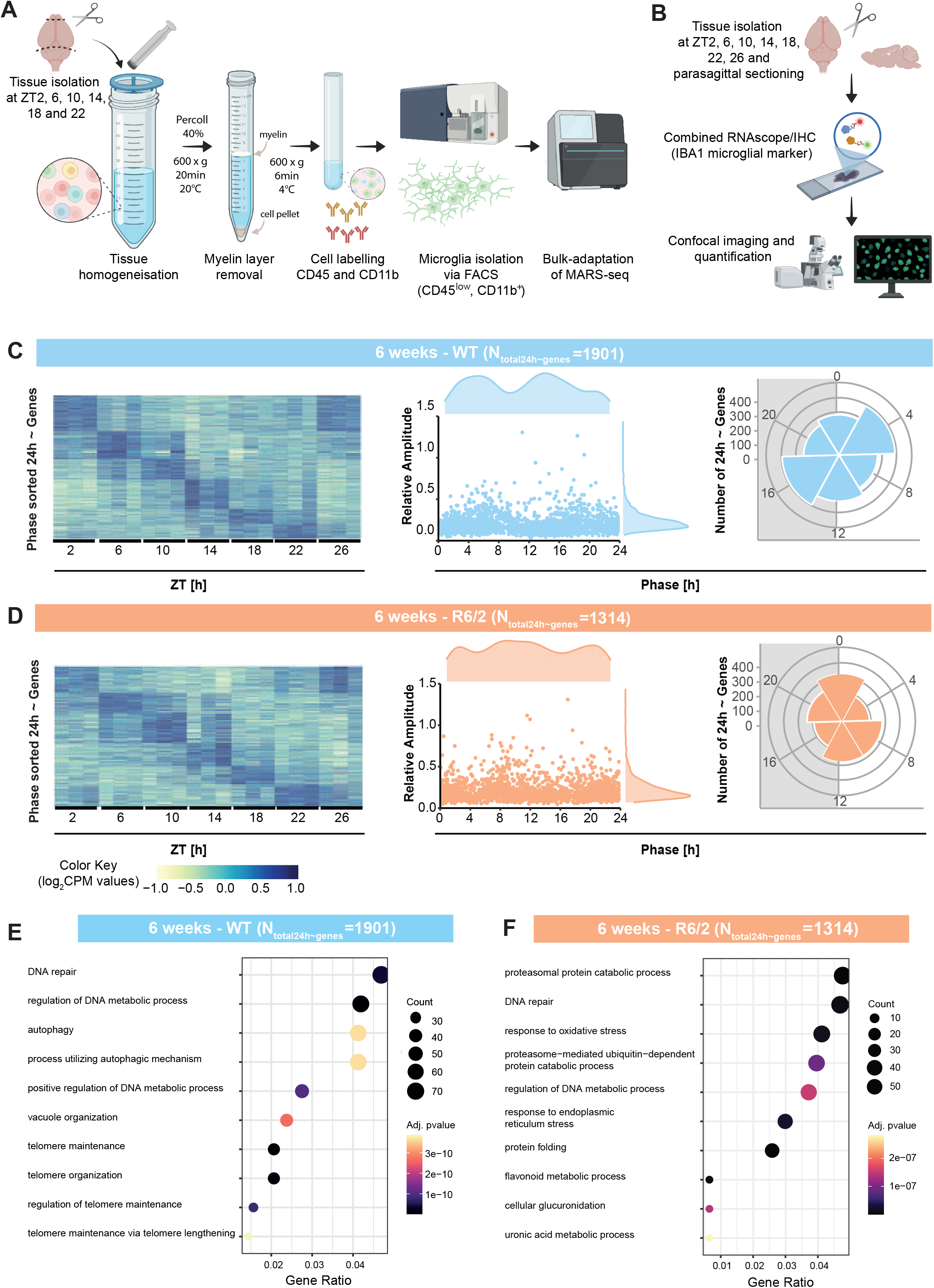
Diurnal gene expression of microglia from 6-week-old WT and R6/2 mice in time-course RNA-seq datasets. (**A**) Experimental workflow for RNA-seq. Mice were kept under a 12:12 LD (light-dark) cycle and fed *ad libitum*. Brains were collected every 4 hours during a time-course of 24 hours. For RNA-seq, dissected brain tissues were homogenized, and microglia were isolated, labelled with CD45/CD11b antibodies, sorted via FACS and subsequently sequenced. (**B**) Experimental workflow for RNAscope hybridization combined with Iba1 immunohistochemistry. (**C**) Phase-sorted heatmaps of significantly 24-hour rhythmic genes with each gene scaled to a range between -1 and 1 with dark blue (1.0) and yellow (- 1.0), respectively, indicating the maximum and minimum gene expression values (left panel); phase and amplitude density plots for 24-hour rhythmic genes (middle panel); Circular plots showing the phase distribution for significantly 24-hour rhythmic genes (q<0.05). White background indicates the light period (lights on) and the grey shaded area indicates the dark period (lights off; right panel) for 6-week-old WT (light blue) and (**D**) R6/2 mice (light orange). (**E**) Top 10 GO (Biological Processes) enriched in 6-week-old WT and (**F**) R6/2 mice.

We next analyzed biological processes in which these circadian expressed genes are involved. At 6 weeks of age, the top 10 significantly enriched GO biological processes in WT microglia included DNA repair, regulation of metabolic processes, telomere regulation and autophagy (**Figure 1E**). Other enriched biological processes were related to proteostasis (**Figure 1E**). For R6/2 microglia at 6 weeks of age, 24-hour rhythmic genes were enriched for biological pathways associated with protein folding, protein degradation (via proteasome or not mediated by a ubiquitin-dependent process), response to oxidative stress and response to endoplasmic reticulum stress, four biological functions that have been reported to be associated with HD^42–44^ (**Figure 1F**). Other biological pathways enriched in microglia from 6-week-old R6/2 mice involved DNA repair, flavonoid metabolic process, cellular glucuronidation and uronic acid metabolic process (**Figure 1F**).

We then investigated diurnal gene expression in microglia from symptomatic 15-week-old R6/2 and age-matched WT mice. We identified 1244 genes as significantly 24-hour rhythmic in WT microglia and 1208 genes as significantly 24-hour rhythmic in R6/2 microglia (**Figures 2A-B**). Our phase-sorted heatmaps showed robustly rhythmic genes in microglia, regardless of the genotype (**Figures 2A-B**). Contrary to the observation for the 6-week-old mice, the phase distribution of most 24-hour rhythmic genes for the 15-week-old mice, regardless of genotype, displayed a bimodal distribution with the majority of the phases grouped at dusk and dawn (**Figures 2A-B**).

**Figure 2.**
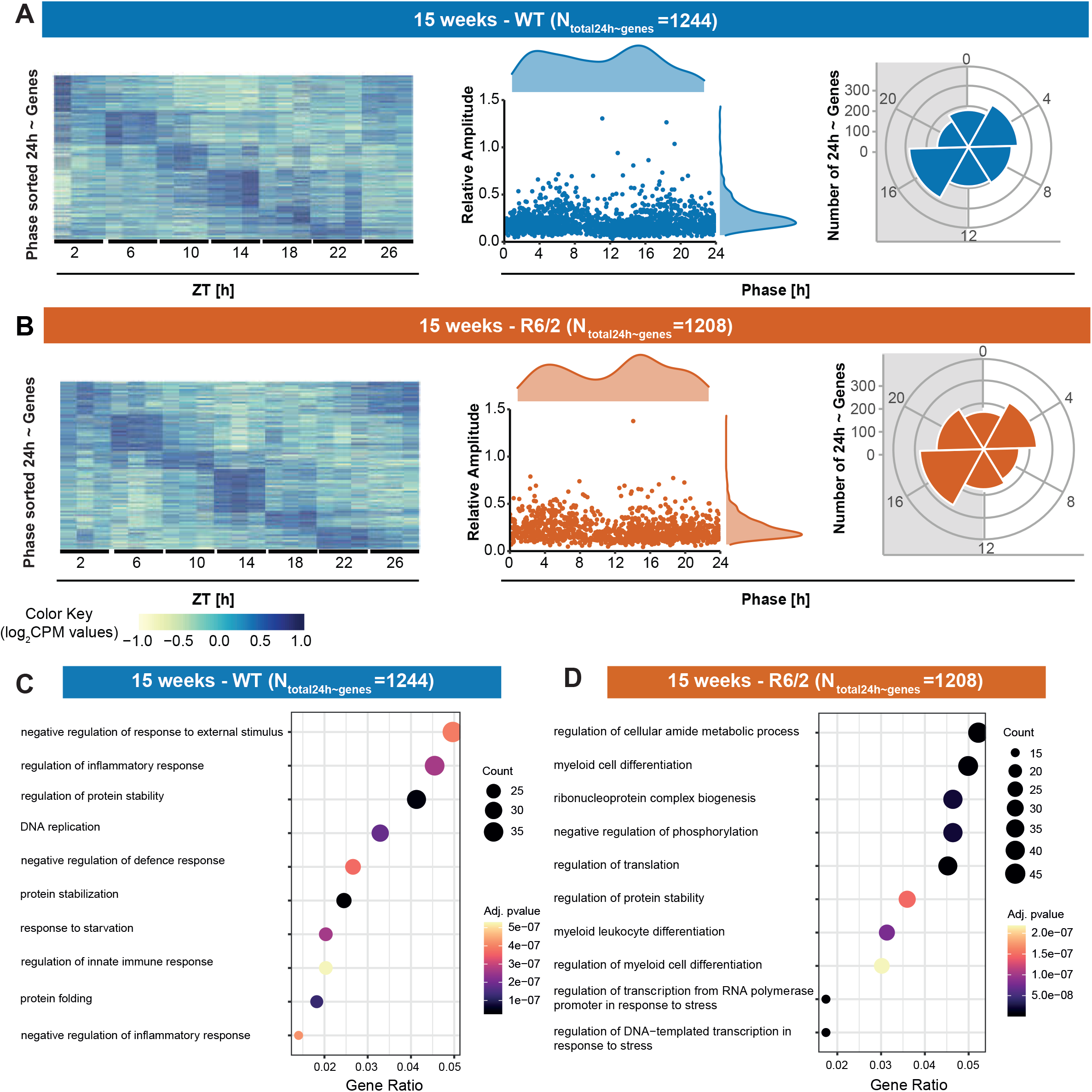
Diurnal gene expression in microglia from 15-week-old WT and R6/2 mice in time-course RNA-seq datasets. (**A**) Phase-sorted heatmaps of significantly 24-hour rhythmic genes (q<0.05) (left panel). Each gene expression value is scaled to a range between -1 and 1 with dark blue (1.0) and yellow (-1.0), respectively, indicating a peak and trough of expression; phase and amplitude density plots for 24-hour rhythmic genes (middle panels); Circular plots show the phase distribution for significantly 24-hour rhythmic genes (q<0.05); white background indicates lights on, and the grey shaded area indicates lights off period (right panel) for 15-week-old WT mice (dark blue) and (**B**) R6/2 mice (dark orange). (**C**) Top 10 GO (Biological Processes) enriched in 15-week-old WT and (**D**) R6/2 mice.

Interestingly, in 15-week-old WT mice, the 24-hour rhythmic genes were associated with biological processes involved in regulation of inflammatory, defence and innate immune responses (**Figure 2C**). Other biological processes enriched were related to protein stability and folding. In 15-week-old R6/2 mice, 24-hour rhythmic genes were enriched in biological processes involved in myeloid cell differentiation and its regulation, protein stabilization, DNA repair, regulation of transcription and translation, and regulation of the cellular amide metabolic process (**Figure 2D**).

The subsequent comparative rhythmicity analysis for 6-week-old WT and R6/2 mice resulted in a total of 2910 circadian rhythmic genes (rhythmic in at least one group). Out of those, 93.7%, (2720 genes) exhibited similar circadian oscillatory patterns, 6.3% (190 genes) were differentially rhythmic (DR), undergoing either a significant change, gain or loss in their 24-hour circadian expression (q<0.05). For the differentially rhythmic genes in 6-week-old mice, we observed a loss of oscillation among R6/2 mice for 41.1% of the genes, and a gain of circadian rhythmicity for 35.8%, while 23.1% of DR genes displayed changes in amplitude and/or phase (**Figure 3A**). In 15-week-old-mice, 89.4% (1854 out of 2074 genes) were significantly 24-hour rhythmic in WT and R6/2 mice. Out of those, 10.6 % (220 genes) underwent a significant change, gain or loss of oscillations with 41.4% of genes gaining rhythmicity and 40.4% showing a loss of circadian rhythms among R6/2 mice, while 18.2% showed a change in amplitude and/or phase (**Figure 3A**).

**Figure 3.**
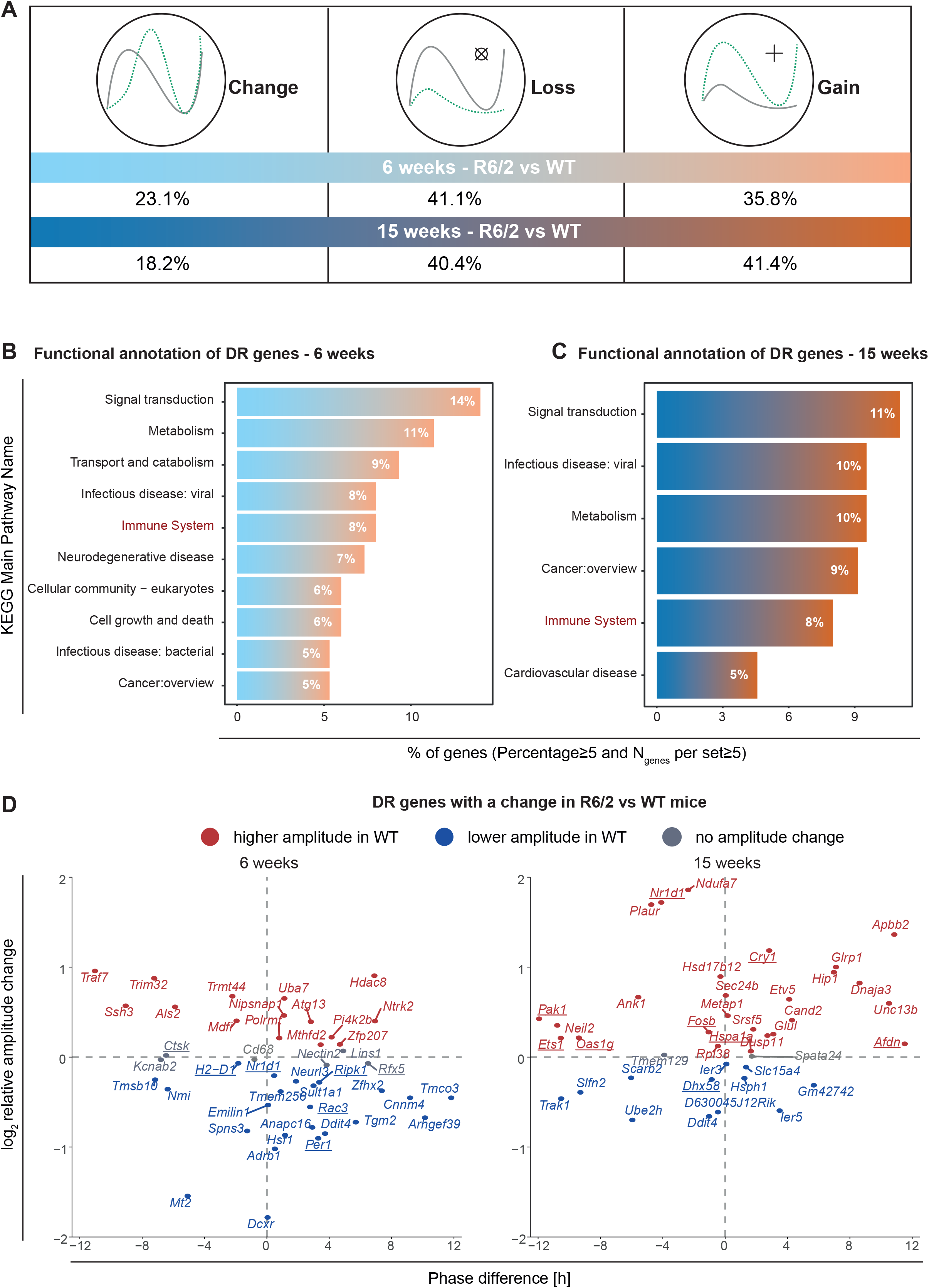
Differentially rhythmic genes in microglia from 6-week-old and 15-week-old R6/2 versus WT mice. **(A)** Changes in rhythmicity (amplitudes and/or phases) of microglial 24-hour rhythmic genes from WT and R6/2 mice (q<0.05). Functional annotation of differentially regulated (DR) genes (q<0.05) in **(B)** 6-week-old and **(C)** 15-week-old mice. **(D)** Genes with a significant differential rhythmic pattern (q<0.05) are colour-coded according to the change in amplitude values. Higher amplitude in WT mice (red), lower amplitude in WT mice (blue) and DR genes without an amplitude change |log_2_| <0.1 (grey). Genes from immune-system pathways and/or core-clock gene networks are underlined.

Subsequently, we performed a functional annotation of differentially rhythmic genes by mapping DR genes to different KEGG pathways. Our results indicate a higher proportion of genes involved in signal transduction, metabolism, transport, and catabolism, followed by pathways related to viral infectious diseases, particularly in 6-week-old mice (**Figure 3B**). Across both age groups, signal transduction emerged as the predominant pathway represented by the highest proportion of genes, while the immune system was another overrepresented pathway in both age groups (**Figures 3B-C**). In 6-week-old mice, DR genes between WT and R6/2 mice were also associated with pathways related to neurodegenerative diseases, cellular growth and death, bacterial infectious diseases, and cancer pathways (**Figure 3B**). Conversely, in 15-week-old mice, overrepresented pathways of DR genes between WT and R6/2 mice included signal transduction, viral infectious diseases, metabolism, cancer, and cardiovascular diseases (**Figure 3C**). We next aimed at characterizing the set of DR genes that displayed alterations in their circadian properties, namely in amplitude and/or phase (**Figure 3D**). In total, 44 genes showed a change in 6-week-old mice; and 40 genes in 15-week-old mice when comparing R6/2 and WT mice. More genes showed a lower amplitude in 6-week-old WT mice, whereas the opposite was observed in 15-week-old WT mice. Interestingly, among the DR genes between WT and R6/2 mice were the core clock genes *Nr1d1* and *Per1,* which exhibited a lower amplitude and phase delay of 1 hour and 3 hours, respectively, in 6-week-old mice (**Figure 3D**). In contrast, in the 15-weeks-group, *Nr1d1* had a higher amplitude and a 4-hour phase advance in WT mice. *Cry1,* another core clock component, also showed an increase in amplitude and a phase delay of ∼3 hours in 15-week-old WT mice (**Figure 3D**).

### Circadian dysregulation of core clock and immune-related genes expression in microglia from symptomatic R6/2 mice

We next analyzed the change in oscillatory properties of 9 core clock genes^45^ (*Arntl*, *Ror*α, *Cry1*, *Cry2*, *Nr1d1*, *Nr1d2*, *Per1*, *Per2*, and *Per3*) expressed in microglia and detected as rhythmic in at least one of the conditions in our RNA-seq dataset (**Figures 4A-B**). At 6 weeks of age, all of the clock genes tested expressed 24-hour rhythms in microglia from WT mice (**Figures 4A-B**). We found similar rhythmic properties for microglia from WT mice at 15 weeks of age, showing an anti-phasic relationship between the positive element, *Arntl*, and the negative elements, *Cry1/2* and *Per1/2/3* (**Figures 4A-B**). *Nr1d1* expression displayed a small phase delay (∼1 hour), whereas *Per1* showed a 3-hour delayed phase in microglia from 6-week-old WT mice (**Figures 4A-B**). In microglia from 6-week-old R6/2 mice, all clock genes tested expressed 24-hour rhythms, except for *Ror*α (**Figures 4A-B**), which showed a significant loss of rhythmicity (**Figure 4B**). The rhythmic profile of *Arntl* was, as expected, in antiphase compared to *Per1* (**Figures 4A-B**) in microglia from 6-week-old WT and R6/2 mice, with phases corresponding to ∼ZT4 and ZT16, respectively. At 15 weeks of age, microglia from R6/2 mice lost rhythmicity in *Arntl* (**Figure 4B**). Most notably, we observed massive loss of rhythmic gene expression in microglia from 15-week-old R6/2 mice with only 4 out of the 9 genes tested showing significant oscillations with phases within ZT12-18 (*Per1*, *Per3*, *Cry1* and *Nr1d1*; **Figures 4A-B**). We observed delayed circadian phase for *Nr1d1* and advanced circadian phase for *Cry1* in microglia from 15-week-old R6/2 mice (**Figure 4B**).

**Figure 4.**
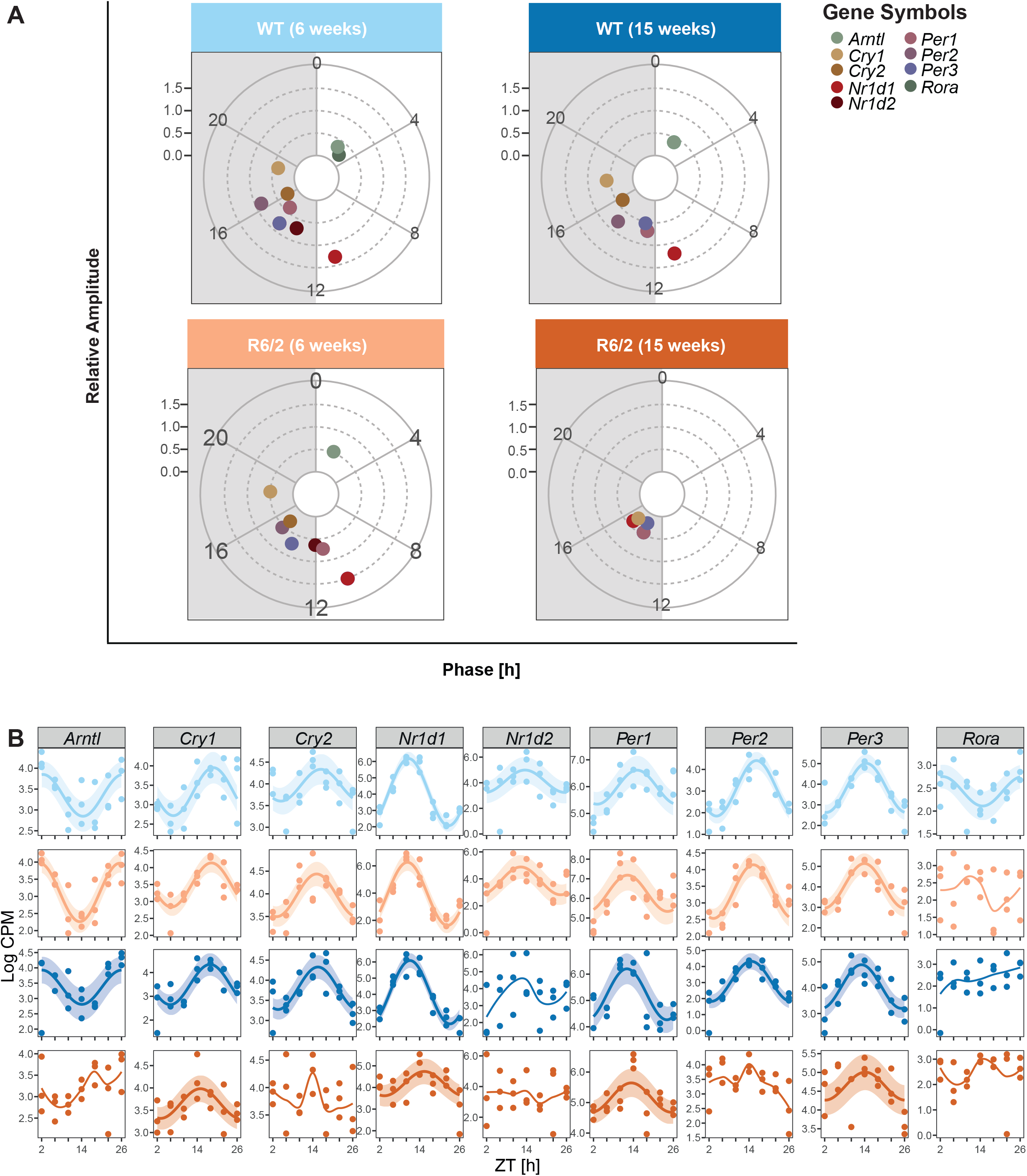
Rhythmicity profiles of core clock network genes vary in microglia from 6-week-old and 15-week-old WT and R6/2 mice. (**A**) The circular plots depict oscillatory properties (relative amplitude and phase distribution) for significantly 24-hour rhythmic core clock genes for: 6-week-old WT mice (light blue), 15-week-old WT mice (dark blue), 6-week-old R6/2 mice (light orange), and 15-week-old R6/2 mice (dark orange). White background indicates lights on and the grey shaded area indicates lights off. (**B**) Circadian expression profiles of core clock network genes in WT and R6/2 mice at 6 and 15 weeks of age. Significantly rhythmic genes (q<0.05) are depicted using a harmonic regression curve with confidence intervals; not significantly rhythmic genes are depicted with LOESS (Locally Estimated Scatterplot Smoothing).

Given the important role of microglia in innate immunity, we next investigated the immune pathways that were rhythmically affected in R6/2 microglia compared to WT microglia. We detected 190 differentially rhythmic genes in microglia from 6-week-old R6/2 versus WT mice (change in rhythmicity, loss or gain of rhythmicity in one genotype compared to the other), and 220 differentially rhythmic genes in microglia from 15-week-old R6/2 versus WT mice (**Tables S1 and S2**). Out of those genes, we then extracted the ones involved in immune pathways (**Table S3**). At the age of 6 weeks, we found 15 differentially rhythmic genes (q<0.05) between presymptomatic R6/2 and WT microglia that were involved in 14 different immune pathways (**Figure 5B; Table S4**). Five immune-related genes had a significant change in circadian profile (*Ctsk*, *H2-D1*, *Rac3*, *Rfx5*, *Ripk1*) (**Figure 3D**), 4 immune-related genes had a gain of rhythmicity in R6/2 microglia (*Card6*, *Ccl6, Mrc2*, *Naip2*) (**Figure 5A**), and 6 immune-related genes lost rhythmicity in R6/2 microglia compared to WT microglia (*Gnb5*, *Pik3c3*, *Plpp3*, *Pstpip1*, *Sos2* and also the core clock element, *Ror*α) (**Figure 5A**). GO enrichment analysis revealed that the differentially expressed immune-related genes in microglia from R6/2 versus WT mice were involved in immune pathways associated with NOD-like, chemokine, and Toll-like receptor signalling pathways, antigen processing and presentation, cellular senescence, natural killer cell mediated cytotoxicity, phagosome, Fc gamma R-mediated phagocytosis, B-cell receptor, Fc epsilon RI signalling pathways, and the cytosolic DNA-sensing, RIG-I-like signalling, Th17 cell differentiation, and T-cell receptor signalling pathways (**Figure 5B**). For microglia from 15-week-old symptomatic R6/2 versus WT mice, two core clock genes, *Arntl* and *Per2*, showed a significant loss of rhythmicity and 23 other immune-relevant genes were differentially regulated (q<0.05) and were found to be associated with 19 different immune pathways (**Figure 5C; Table S5**). Seven of these genes showed a significant change in circadian expression (*Afdn*, *Dhx58*, *Ets1*, *Fosb*, *Hspa1a*, *Oas1g*, *Pak1*), 7 genes showed a gain of rhythmicity in R6/2 microglia (*Cdkn2b*, *Icam2*, *Irf7*, *Itgam*, *Msn*, *Polr3gl*, *Vasp*), and 9 genes lost rhythmicity compared to WT microglia (*Adar, Chek2*, *H2-Aa*, *Irf9*, *Jag2*, *Sec61g*, *Stat2*, *Sting1*, *Tgfbr1*) (**Figure 5C**). The 6 new immune pathways affected in microglia from 15-week-old symptomatic R6/2 mice compared to the 6-week-old R6/2 mice involved signalling pathways of IL-17, C-type lectin receptor, Th1 and Th2 cell differentiation, hematopoietic cell lineage, leukocyte transendothelial migration, and platelet activation based on GO enrichment analysis (**Figure 5D**).

**Figure 5.**
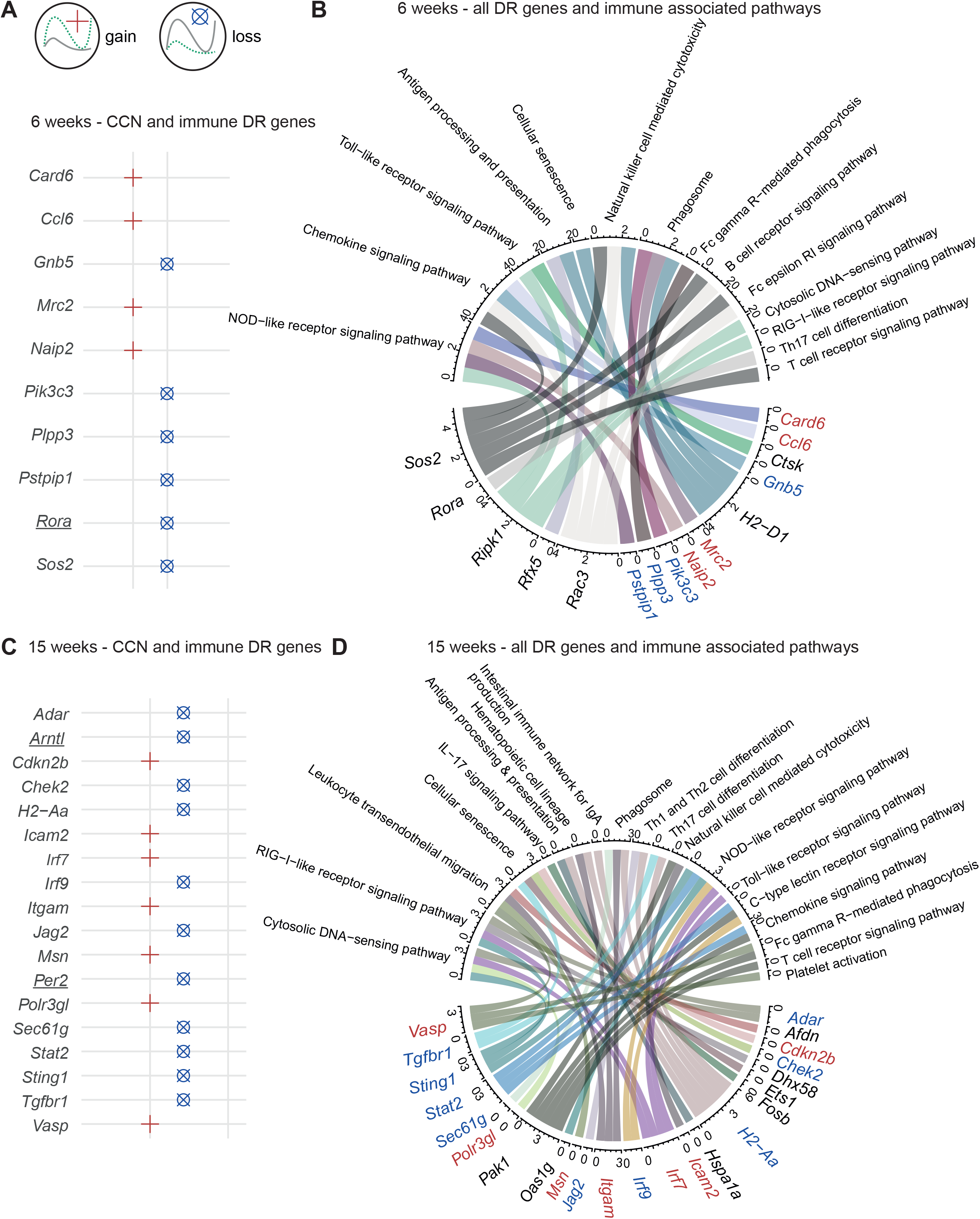
Core clock and immune-associated genes with change, loss or gain of 24-hour rhythmicity in microglia from R6/2 mice. Core clock and immune-associated genes displaying a significant change, gain or loss of their 24-hour rhythmicity (q<0.05) in microglia from R6/2 mice compared to WT mice are shown for (**A**) 6-week-old and (**C**) 15-week-old groups. The chord diagrams represent the differentially rhythmic immune-related genes (loss in red, gain in blue and change in black) and their associated GO enrichment analysis pathways in (**B**) 6-week-old and (**D**) 15-week-old mice.

### Microglia display significant differences in 24-hour gene expression across brain regions in health and disease

We next assessed regional diurnal profiles of gene expression in individual microglia from the two brain structures most affected by HD, namely striatum and cortex^41^, for candidate genes based on the RNA-seq analysis. We selected the core clock gene, *Arntl,* and the immune-related genes, *Ccl2* and *Irf7*, for further analysis by RNAscope based on their circadian expression profiles and differential regulation between WT and R6/2 mice. The core clock gene, *Arntl*, also plays a role in the immune system: *Arntl* directly inhibits the expression of *Ccl2* and indirectly inhibits the expression of *IL-6* via *Nr1d1*^46^. *Ccl2* is upregulated in the striatum in neurodegenerative diseases such as AD and HD^47, 48^ compared to controls. Interferon regulatory factor 7 (*Irf7)* is differentially expressed in an *in vitro* model of HD^40^.

We performed RNAscope combined with Iba1 immunohistochemistry and morphological identification of microglia on parasagittal brain sections from WT and R6/2 mice at 15 weeks of age collected at ZT2, 6, 10, 14, 18 and 22, focusing on striatum and cortex (**Figure 1B**; **Figure 6B** for striatal examples at ZT6 and ZT18; **Supplementary Video** for example of 3D-reconstructed microglia with Imaris software (Bitplane)). Daily profiles for RNAscope data sets for *Arntl*, *Irf7* and *Ccl2* mRNA expression in microglia from 15-week-old WT and R6/2 mice are shown in **Figure 6A** for striatum and in **Figure 6C** for cortex, indicating substantial variations between individual microglia from the same and different brain regions at a given time of day (significance was reached for *Arntl* mRNA expression in microglia from WT mice comparing cortex to striatum as shown in **Table S8**). Notably, 24-hour rhythmic expression of *Arntl* was conserved in striatal microglia from symptomatic R6/2 mice compared to WT mice (Rsq = 0.51 and 0.56) (**Figure 6A; Table S6**). However, we detected non-significant 24-hour rhythmicity for microglial *Arntl* gene expression in the cortex of 15-week-old R6/2 mice (Rsq = 0.34) compared to WT mice (Rsq =0.64) (**Figure 6C; Tables S6 and S7**). As for the immune-related genes, we found that the expression of *Ccl2* and *Irf7* genes in microglia were not significantly 24-hour rhythmic in cortical and striatal microglia from WT mice (**Figures 6A and 6C**). In contrast, cortical microglia from symptomatic R6/2 mice *de novo* acquired 24-hour rhythmic expression of *Ccl2* and *Irf7* genes at the single-cell level, which was not the case for striatum and did not reach statistical significance (**Figures 6A and 6C; Tables S6 and S8**). The gain of oscillation of *Irf7* is in line with our previous analysis using RNA-seq data (**Table S8**).

**Figure 6.**
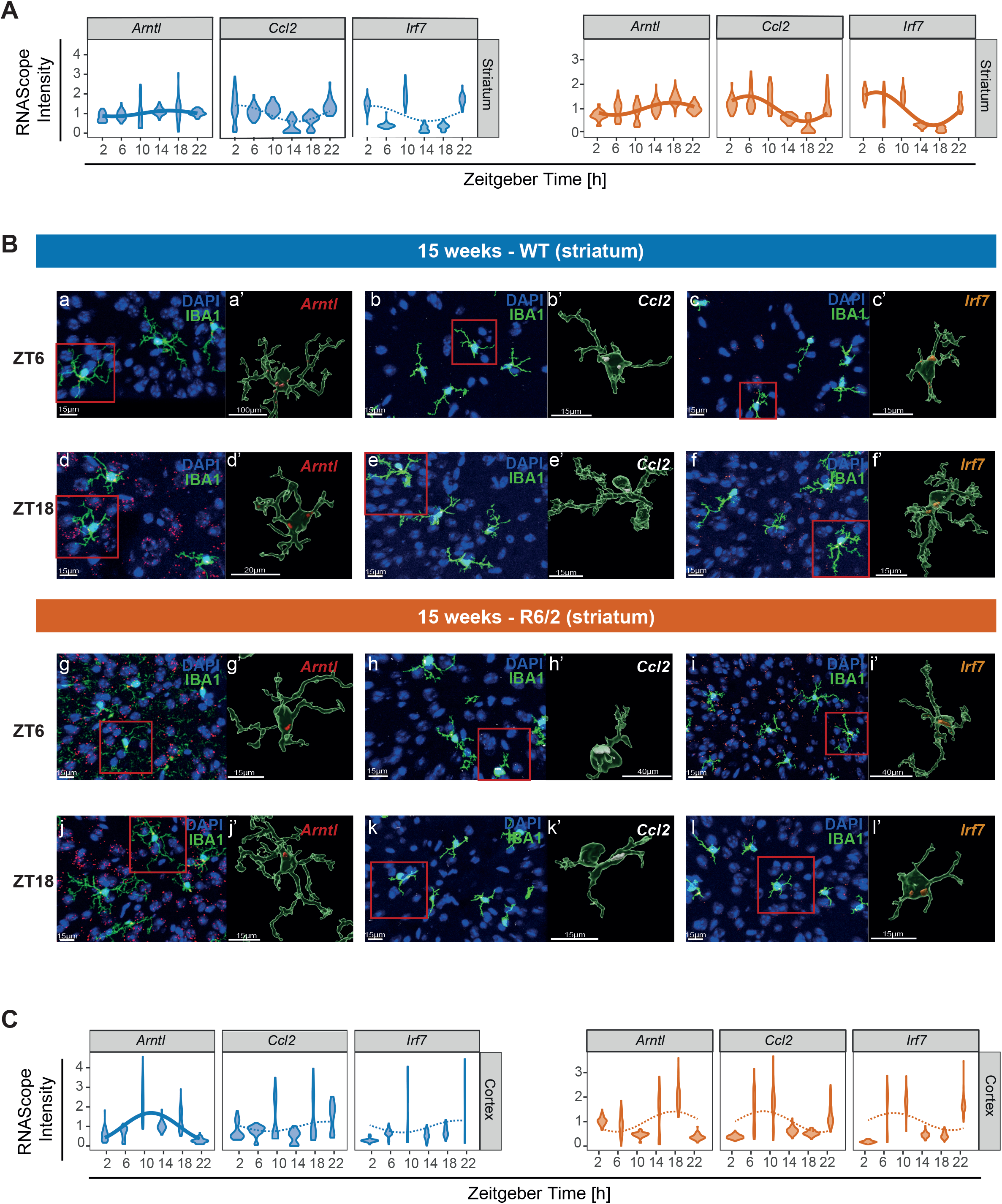
Microglial *Arntl*, *CCl2* and *Irf7* 24-hour gene expression profiles at the single-cell level in 15-week-old WT and R6/2 mice. **(A)** Diurnal mRNA expression profiles of *Arntl*, *Ccl2* and *Irf7* in striatum of 15-week-old WT and R6/2 mice obtained with RNAscope probes, shown with violin plots. Full lines represent significantly 24-hour rhythmic genes (R^2^>0.5) while disconnected lines represent non-significant genes. **(B)** Representative confocal microscopic images (a, c, e, g, i, k) of striatum from 15-week-old WT and R6/2 mice immunostained for IBA1 (marker for microglia; green), 4’,6-diamidino-2-phenylindole (DAPI; blue) for nuclear identification, and hybridized with RNAscope probes for *Arntl* mRNA(red), *Irf7* mRNA (orange) or *Ccl2* mRNA (white). The images in b, d, f, h, j and i depict 3D-rendered microglia with RNAscope signal, magnified from the rectangle indicated in the corresponding confocal images to the left. **(C)** Diurnal mRNA expression profiles of *Arntl*, *Ccl2* and *Irf7* in cortex from 15-week-old WT and R6/2 mice via RNAscope are shown with violin plots. Full lines represent significantly 24-hour rhythmic genes (R^2^>0.5), while disconnected lines represent non-significant genes.

## Discussion

The immune response is one of the many physiological functions that are controlled by the circadian clock. Microglia are the tissue-resident macrophages of the CNS and responsible for the immune surveillance and defence of the CNS. Since the demonstration of the expression of clock genes and the existence of an intrinsic molecular clock in microglia^16, 17^, the circadian regulation of microglial immune response has become an area of active research. Studies in mice revealed important regulation of immune responses in microglia by the core clock proteins ARNTL, CLOCK and NR1D1: for example, the protein heterodimer formed by ARNTL:CLOCK regulates both TLR9^49^ and CCL2 expression^12^ while NR1D1 inhibits the expression of IL-6 by microglia^12^. Accordingly, the response of microglia to immune stimulation by activation of TLR4 via bacterial endotoxin lipopolysaccharide (LPS) is time-of-day dependent^46^ via an *Arntl*-dependent increase in IL-6 expression^21^. As a result, rodents are more prone to sepsis when challenged with LPS during their resting phase^50, 51^.

Since the circadian time aspect seems critical for microglial functions, we aimed to gain further knowledge on the role of the microglial circadian clock in WT mice and the R6/2 transgenic mouse model of HD. Our RNA-seq data revealed robust 24-hour rhythmic gene expression in healthy microglia, enriched at 6 weeks of age in DNA repair and metabolic processes, telomere regulation, and autophagy, whereas at 15 weeks of age pathways involved in protein stability/folding, regulation of inflammatory, defence and innate immune response were enriched. Microglia are heterogeneous cells whose states vary depending on the signals they receive from their environment which change with age^52^. The differences we see between microglia from 6-week-old and 15-week-old WT animals may reflect these ongoing changes in microglial functions. The 9 core clock genes tested (*Arntl*, *Ror*α, *Cry1*, *Cry2*, *Nr1d1*, *Nr1d2*, *Per1*, *Per2*, and *Per3*) expressed 24-hour rhythms in WT mice, with anti-phasic rhythms between the positive element, *Arntl*, and the negative elements, *Cry1/2* and *Per1/2/3* genes. Interestingly, at 15 weeks of age, *Ror*α and *Nr1d2* lost 24h-hour rhythmic gene expression in WT microglia.

Microglia play a key role in ensuring an optimal milieu for neuronal activity and function^53^. Consequently, glial cell dysregulation may contribute to the pathogenesis of neurodegenerative diseases, such as HD^54^. In fact, early disturbances of the circadian clock and immune system activation have been described prior to the onset of motor and cognitive symptoms in HD. Glial cells expressing mHTT contribute to neuronal excitotoxicity in R6/2 mice ^55^. Mutant HTT is highly expressed in immune cells and while the depletion of mHTT from microglia did not rescue the phenotype of BACHD transgenic mice^56^, several lines of evidence suggest that microglial mHTT expression promotes cell-autonomous inflammatory immune responses with increased expression and transcriptional activity of PU.1 and CCAT/enhancer-binding protein^35^ and transcriptional activation of pro-inflammatory genes in HD^57–59^. As a result, the prolonged activation of microglia and their consecutive production of inflammatory mediators may contribute to neurodegeneration in HD^36^.

Our results suggest that mHTT has a deleterious effect on the microglial circadian clock. It is noteworthy that neither the expression of *Htt* nor the genes encoding the interacting proteins, *Hip1* (Huntingtin-interacting protein 1) or *Hap1 (*Huntingtin-associated protein 1), were found to be 24-hour rhythmic in our RNA-seq analysis. When we assessed the daily gene expression rhythms in microglia from R6/2 mice, we found that circadian profiles of specific core clock genes were particularly affected. Although the expression patterns of all clock genes tested (with the exception of *Ror*α) were 24-hour rhythmic in presymptomatic R6/2 mice, 5 core clock genes lost 24-hour rhythmicity at the symptomatic stage (*Arntl*, *Ror*α, *Cry2*, *Nr1d2* and *Per2*). The 24-hour rhythmic gene expression profile of *Arntl* was, as expected, in anti-phase compared to *Per1*, but microglia from symptomatic R6/2 mice lost 24-hour rhythmicity of *Arntl* gene expression. Only 4 genes belonging to the negative feedback loop (*Per3*, *Per1*, *Nr1d1* and *Cry1*) retained 24-hour rhythmicity of gene expression in microglia, raising questions as to how this partial preservation may impact on overall disease progression and immune responses. Our RNAscope results suggest that there are substantial differences in phases and profiles of *Arntl* expression at the single-cell level for microglia within the same brain region and between cortex and striatum for a given time of the day. Such intra-tissue desynchrony has previously been described for peripheral tissues and is increasingly recognized in the circadian field^58^. Methodological differences may explain the discrepancy of our results with recently published data suggesting global preservation of diurnal gene co-expression modules between cortex and striatum^60^.

Besides the changes in diurnal clock gene expression in microglia from R6/2 mice, we also observed loss of 24-hour rhythmicity in the expression of immune-relevant genes, including *Adar*, *Chek2*, *H2-Aa*, *Irf9*, *Jag2*, *Sec61g*, *Stat2*, *Sting1*, *Tgfbr1*. These changes may have implications for HD pathogenesis, e.g., Stat2 is part of the JAK/STAT signalling pathway which is disrupted in HD. Notably, we also detected *de novo* 24-hour rhythmicity in the expression of immune-relevant genes, such as *Cdkn2b*, *Icam2*, *Irf7*, *Itgam*, *Msn*, *Polr3gl* and *Vasp*, in microglia from symptomatic R6/2 mice. *Irf7* is a transcription factor involved in the Toll-like receptor signalling pathway and was found to be differentially expressed in a *mHTT* knock-in cell culture model of HD^40^ and in our own RNA-seq analysis of peripheral blood cells obtained from HD patients^61^. The reason behind this emergence of rhythmic expression of immune-relevant genes in microglia in HD requires further exploration.

Our findings of circadian reprogramming of microglia in the context of neurodegeneration are intriguing and require confirmation in other neurodegenerative diseases, e.g., Alzheimer’s disease. It is tempting to speculate that modulating microglial function in the context of circadian immune therapy may be a promising avenue for future therapeutic interventions for neurodegenerative diseases.

## Acknowledgements

We would like to thank Tanja Specowius, Jasmin Jamal El-Din and Christian Böttcher for their excellent technical support and Bert Maier as well as Camila Fernández Zapata for helpful technical discussion. We would also like to thank BCRT Flow Cytometry Lab for their support in sorting microglia for the bulk RNA sequencing study. Icons from panels A and B of Figure 1 were retrieved from BioRender (full license). This work was funded by the German Research Foundation (SFB/TRR167/2 B07) to JP. The work in the AR group was funded by the MSH Medical School Hamburg and by the Dr. Rolf Schwiete Stiftung.

## Author Contributions

Conceptualization, J.P.; Methodology, K.O., M.Y., J.P., A.R.; Investigation, K.O., M.Y., L.R., H.S.K., E.Y., A.Kn., C.B., A.Kr., I.A., A.R., J.P.; Formal Analysis, M.Y. and E.Y., Writing – Original Draft, K.O.; Writing – Review & Editing, K.O., M.Y., J.P., A.R.; Visualisation, K.O. and M.Y.; Resources, J.P., A.Kr., A.R.; Supervision, J.P. and A.R.; Funding acquisition, J.P. and A.R.

## Methods

All experimental procedures were performed in accordance with the GV-SOLAS and FELASA guidelines of the animal care of the Charité – Universitätsmedizin Berlin and with the approval of the local federal state authorities “Landesamt für Gesundheit und Soziales” (LAGeSO, Berlin, Germany) under the license numbers T0157/11 and G0031/21.

### Animal model

The mouse line used in this study was obtained from the Jackson Laboratory: B6CBA-Tg(HDexon1)62Gpb/1J; Strain #:002810; RRID:IMSR_JAX:002810; common name: B6CBA-R6/2. R6/2 and WT female mice used for the study were generated in the breeding facility in FEM-Bayer (Berlin) on a CBA × C57BL/6J background (total of n=42 for RNA-seq and n=36 for RNAscope). Mice were then housed in specific pathogen-free (SPF) animal facilities of the Charité – Universitätsmedizin Berlin. R6/2 transgenic mice ubiquitously express the exon 1 of human Huntingtin (3% of the N-terminal region of the protein that incorporates the polyglutamine expansion^62^ with a CAG repeat expansion). Genotype identification was conducted on ear snips and verified after completion of the experiments, using the forward primer 5’-CCGCTCAGGTTCTGCTTTTA-3’ and the reverse primer 5’-GAGTCCCTCAAGTCCTTCCA -3’. The determination of the number of CAG repeats was verified by Laragen (Los Angeles, USA). In this study, we used R6/2 mice and chose to test them at two different stages of disease: presymptomatic (6 weeks) and symptomatic (15 weeks) to evaluate whether circadian gene expression in HD microglia is disrupted. For the experiments, mice were group-housed in cages of a maximum of 6 animals but mixed genotype. For habituation and entrainment to 12:12 LD cycle, mice were group-housed in a ventilated, light-tight and sound-proof Scantainer cabinet (Scanbur, Denmark), with a built-in light system (maximum 300 lux) and controlled humidity and temperature, during 10-14 days prior to brain tissue collection. Mice had *ad libitum* access to food and water. Tissue was then collected in a time-course manner to identify and analyze 24-hour rhythmic genes (**Figure 1A**).

Mouse brains were collected from R6/2 mice at 6 weeks and 15 weeks of age and were compared with age-matched WT littermates, at targeted time of day (ZT2, 6, 10, 14, 18, 22 or 26, with ZT0 corresponding to lights switched on and ZT12 with lights switched off) with n=3 for each time point and genotype. This sample size was decided after statistical consultation with the Institute of Biometry and Clinical Epidemiology at Charité (Berlin). Brains were collected after anaesthesia with 4 % isoflurane and after checking that there was no reaction to the interdigital reflex to ensure that the animal has reached a surgical level of anaesthesia.

### RNA sequencing

Brain tissue was immediately collected in cold 1xPBS, then homogenized by gentle mechanical dissociation using syringe plunger and strainer. Microglia were subsequently enriched by 40% Percoll gradient. After washes with FACS buffer, cells were blocked with Fc Block (1:100; Biolegend ref 101320) and subsequently stained with CD45-FITC (1:100; ref 101212: Biolegend) and CD11b-APC (1:100; ref 103108 Biolegend) for final purification by flow cytometry (Supplementary Figure 1 for gating strategy) using a FACS Aria-II cell sorter (BCRT, Berlin, Germany). The microglia were directly sorted into a lysis buffer (Invitrogen) from the Dynabeads™ mRNA DIRECT™ Purification Kit (Invitrogen). Samples were immediately frozen on dry ice and then stored at -80°C prior to RNA-seq analysis.

The RNA extraction was performed using the Dynabeads™ mRNA DIRECT™ Purification Kit (Invitrogen) according to the manufacturer’s protocol. A total of 10^4^ cells from the microglia population was sorted into 40-200 µL of lysis/binding buffer (Life Technologies). mRNA was captured with 12 µl of Dynabeads oligo(dT) (Life Technologies), washed, and eluted at 85°C with 6.5 ml of 10 mM Tris-Cl (pH 7.5). A bulk adaptation of the MARS-Seq protocol^63, 64^ was used to generate RNA-seq libraries for expression profiling of microglia from R6/2 mice and control. Briefly, RNA from each sample was barcoded during reverse transcription and pooled. Following Agencourt Ampure XP beads cleanup (Beckman Coulter), the pooled samples underwent a second strand synthesis and were linearly amplified by T7 in vitro transcription. The resulting RNA was fragmented and converted into a sequencing-ready library by tagging the samples with Illumina sequences during ligation, reverse transcription, and PCR. Libraries were quantified by Qubit and TapeStation as well as by qPCR for *ActB* gene as previously described ^64^. Sequencing was done on a Nextseq 75 cycles high output kit (Illumina; paired end sequencing with gene read from one side of the library and the barcode read from the other side, but with genes only analyzed in a single read mode).

### RNA-seq data processing

Quality control of raw reads was assessed using FastQC (version 0.11.5). Reads were demultiplexed to extract sample information from the barcodes and aligned to the mouse genome (Mus musculus, GRCm38/mm10) using HISAT (v.0.1.5). Multiple mapping reads were excluded. Duplicate reads that consist of identical UMIs were removed. Reads were subsequently quantified; expression levels were determined at the exon level using HOMER software (http://homer.salk.edu) and corresponding genes were annotated using USCS Genome Browser. Raw counts were normalized using the TMM (weighted trimmed mean of M-values) method from R package edgeR (v3.20.9) and subjected to log transformation. A low gene expression threshold was used based on the mean expression levels over the entire 24-hour time series, genes with ≥ 0.5 CPM were retained and counts were then renormalized.

### RNAscope and immunohistochemistry

Mouse brains (n =3 females for each time point and genotype) from R6/2 mice at 6 weeks and 15 weeks of age and from age-matched WT littermates were quickly harvested (within 5 min), at targeted time of day (ZT2, 6, 10, 14, 18 and 22) directly immerged in PFA 4% for 2 days and cryoprotected in a sucrose gradient for 3-4 days. Brains were then flash frozen using isopentane cooled at -35°C in dry ice, then stored at -80°C until sectioning. Parasagittal brain sections at 16µm thickness were obtained using a cryostat set at -20°C. Brain sections were directly mounted on SuperFrost® Slides (Fisher Scientific, Cat# 12-550-15), allowed to dry at room temperature for 1 hour and stored at -80°C until performing the RNAscope assay for simultaneous detection of 2 RNAscope probes.

For this, we used the RNAscope Multiplex fluorescent V2 Assay (Cat# 323100; ACDbio, Biotechne, Wiesbaden-Nordenstadt, Germany) following the manufacturer’s guideline. Briefly, sections were immerged for 25 minutes in PFA 4% on normal crushed ice, then dehydrated in a gradient of ethanol from 50% to 70% then 100%. After air drying and creating a hydrophobic barrier (ImmEdge pen), the sections were incubated with H_2_O_2_ for 10 min at RT, washed, and incubated with protease IV for 25 minutes. After two washes, sections were finally hybridized with the RNAscope probes for *Ccl2*, *Arntl* or *Irf7* (Cat# 311791, 438741-C2, 534541-C3; ACDbio, Biotechne, Wiesbaden-Nordenstadt, Germany) for 2 hours at 40°C. We hybridised the probes for *ccl2* (channel 1), *irf7* (channel 3) on the same sections as they were customised on different channels. At this stage, sections were then stored in 5x SSC buffer overnight until resuming the protocol. For the amplification step, sections were subsequently incubated with AMP1 (channel for gene 1) and AMP2 (channel for gene 2) for 30 min at 40°C with intermittent washes. For fluorescent detection of gene 1, sections were then incubated with HRP-channel of gene 1 for 15 min at 40°C and Opal 620 (1:1500; Akoya, Biosciences, Cat# FP1495001KT) diluted in TSA buffer (Akoya, Biosciences) or Opal 690 (1:1500; Akoya, Biosciences, Cat# FP1497001KT) for 30 min at 40°C, then washed and blocked with RNAscope® Multiplex FL v2 HRP blocker. For fluorescent detection of gene 2, the sections were then incubated with HRP-channel of gene 2 and Opal 620 or Opal 690, using the Opal different from gene 1, and finally incubated with the HRP Blocker.

After this, sections were washed for immunohistochemistry in TBS 0.5M for 2×2 min, blocked with TBS-Triton 0.3% for 10 min and blocked in NDS 10% for one hour. Sections were then incubated with goat anti-IBA1 (1:400; Abcam, ab178846) overnight at 4°C. After 2×10min washes in TBS, sections were finally incubated in the secondary antibody Alexa 488 (1:250; Invitrogen, A21206) for 2 hours, washed 2×10 min then nuclei were counterstained using DAPI (1:10000) for 5 minutes. Sections were finally washed and coverslipped, left to dry at RT for a minimum of 16 hours and stored in an air-proof container at 4°C until image acquisition.

### Image acquisition and gene/protein quantification

Images of the brain sections hybridized with two RNAscope probes and immunostained with anti-IBA1 were acquired using a confocal laser-scanning microscope (Leica TCS SP5, Leica Microsystems) at 400x magnification using LAS X software Z stack images were acquired in cortex, hippocampus, striatum and thalamus (3 images per brain region) for all the sections. Images were then analysed with a custom-made protocol using the imaging software Imaris (Bitplane) for visualization of the 3D confocal images and for detection, quantification and co-localization of RNAscope probe signals with microglia (Iba1 channel). For each image, the Iba1 channel was first processed with a Gaussian filter of 0.5 µm and a background subtraction of 20% to reduce background noise. A surface for Iba1 was subsequently created with a surface detail of 0,3 µm. We applied a threshold cut-off when necessary to remove background signal, and the same threshold was always applied for a gene for each picture and across brain regions. We obtained the cell volume for each individual cell and the intensity sum signal for the targeted gene. We then normalized our single-cell data to the cell volume in order to obtain the intensity sum of the RNAscope signal for each gene at the single-cell level.

### Quantification and statistical analysis

#### Rhythmicity analysis

For bulk RNA-seq and RNAscope data, gene expression rhythmicity was assessed using the RAIN algorithm^65^. For the RNA-seq data set (Gene Expression Omnibus (GEO) Accession No: GSE146705), unlogged expression values were used as input and the resulting RAIN p-values were corrected for multiple testing using the Benjamini-Hochberg (BH) method. Statistical significance was assessed based on the q-value threshold for RNA-seq datasets (q-value < 0.05). For RNAscope datasets, to be able to handle differences in number of single cells in different conditions, R-Squared (R^2^) values were used for evaluation of the goodness of the fit and for denoting the significance of the rhythmicity (R^2^>0.5). Circadian-related properties (phase and amplitude) were determined by setting a fixed period of 24 hours and using the harmonic regression (a.k.a. cosinor) method as implemented in the R package DiscoRhythm^66^ fitting a sinusoidal wave to expression values. Absolute amplitude of the 24-hour significant oscillations (the distance between maximum and minimum expression values for RNA-seq dataset and/or maximum and minimum ratio of RNAscope signal per volume of microglia were assessed as A= √(a^2+b^2 ) and phases reflecting the maximum expression time point (peak) formulated as tan φ = b/a. The relative amplitude is determined by division of absolute amplitude value (A) to the mean level (m) as rel_amplitude_=A/m. The Detection of Differential Rhythmicity (DODR) algorithm^67^ was then used to determine the differentially rhythmic patterns by carrying out a pairwise comparison between R6/2 vs WT mice among the same age groups. Statistical significance threshold for genes exhibiting a differential rhythmicity assessed based on BH-adjusted DODR p-values (q-value < 0.05).

### Functional annotation of differentially rhythmic genes

The RefSeq mRNA IDs of the differentially rhythmic genes were used as the input for DAVID gene functional classification tool^68^ and resulting KEGG_Pathway lists were downloaded. The genes and associated pathways were then mapped to the higher hierarchical category to be represented with their global function such as ‘Immune system’ by gathering the list of pathways sub-listed under these modules and removing duplicates. The genes which were represented by at least 5 genes per main pathway category were depicted subsequently in the plots (if they represented ≥ 5% of the differentially regulated genes).

### Identification of the immune-relevant genes with differential rhythmicity between genotypes

The immune-relevant genes shown in **Table S1** (6-week-old WT and R/2 mice) and **Table S2** (15-week-old WT and R/2 mice) were identified using these criteria: from the differentially rhythmic genes which were identified based on the genes which were rhythmic in at least in one of the genotypes per each age group. If a gene was significantly rhythmic in both genotypes but showed a different rhythmicity profile (phase/amplitude), this is indicated as “change” in the tables. If a gene was rhythmic in WT but not in R6/2 mice, this is indicated as “loss”. If a gene was significant in R6/2 but not in the WT mice, this is indicated as “gain”. Next, the genes involved in the KEGG data based immune pathways (mouse module, accession date: 13.06.2023) were extracted (**Table S3**), duplicated genes were removed and intersected with the differentially rhythmic genes identified (**Tables S4 and S5**).

### Data and code availability

- RNA-seq data were deposited in the GEO (Gene Expression Omnibus) repository and will be publicly available upon publication of the manuscript (GEO Accession No: GSE146705, also listed in the key resources table).
- This study did not generate new code. All computational tools, packages and functions used in this manuscript are freely available and are referred to in the corresponding Star Method sections.
- Any additional information required to re-analyze the data reported in this paper is available from the lead contact upon request

## References

1. Ginhoux, F., et al. Fate mapping analysis reveals that adult microglia derive from primitive macrophages. Science 330, 841–845 (2010).

2. Wolf, S.A., Boddeke, H.W. & Kettenmann, H. Microglia in Physiology and Disease. Annu Rev Physiol 79, 619–643 (2017).

3. Borst, K., Dumas, A.A. & Prinz, M. Microglia: Immune and non-immune functions. Immunity 54, 2194–2208 (2021).

4. Prinz, M., Jung, S. & Priller, J. Microglia Biology: One Century of Evolving Concepts. Cell 179, 292–311 (2019).

5. De Luca, S.N., et al. Conditional microglial depletion in rats leads to reversible anorexia and weight loss by disrupting gustatory circuitry. Brain Behav Immun 77, 77–91 (2019).

6. DeBruyne, J.P., Weaver, D.R. & Reppert, S.M. CLOCK and NPAS2 have overlapping roles in the suprachiasmatic circadian clock. Nat Neurosci 10, 543–545 (2007).

7. Cedernaes, J., Waldeck, N. & Bass, J. Neurogenetic basis for circadian regulation of metabolism by the hypothalamus. Gene Dev 33, 1136–1158 (2019).

8. Castaneda, T.R., de Prado, B.M., Prieto, D. & Mora, F. Circadian rhythms of dopamine, glutamate and GABA in the striatum and nucleus accumbens of the awake rat: modulation by light. J Pineal Res 36, 177–185 (2004).

9. Clark, G.T., et al. Circadian control of heparan sulfate levels times phagocytosis of amyloid beta aggregates. PLoS Genet 18, e1009994 (2022).

10. Yu, X.F., et al. T17 Cell Differentiation Is Regulated by the Circadian Clock. Science 342, 727–730 (2013).

11. Scheiermann, C., et al. Adrenergic Nerves Govern Circadian Leukocyte Recruitment to Tissues. Immunity 37, 290–301 (2012).

12. Gibbs, J.E., et al. The nuclear receptor REV-ERBα mediates circadian regulation of innate immunity through selective regulation of inflammatory cytokines. P Natl Acad Sci USA 109, 582–587 (2012).

13. Bellet, M.M., et al. Circadian clock regulates the host response to. P Natl Acad Sci USA 110, 9897–9902 (2013).

14. Brancaccio, M., Patton, A.P., Chesham, J.E., Maywood, E.S. & Hastings, M.H. Astrocytes Control Circadian Timekeeping in the Suprachiasmatic Nucleus via Glutamatergic Signaling. Neuron 93, 1420–1435 e1425 (2017).

15. Tso, C.F., et al. Astrocytes Regulate Daily Rhythms in the Suprachiasmatic Nucleus and Behavior. Curr Biol 27, 1055–1061 (2017).

16. Nakazato, R., et al. Selective upregulation of Per1 mRNA expression by ATP through activation of P2X7 purinergic receptors expressed in microglial cells. J Pharmacol Sci 116, 350–361 (2011).

17. Hayashi, Y., et al. The intrinsic microglial molecular clock controls synaptic strength via the circadian expression of cathepsin S. Sci Rep 3, 2744 (2013).

18. Wang, X.L., et al. Microglia-specific knock-down of Bmal1 improves memory and protects mice from high fat diet-induced obesity. Mol Psychiatry 26, 6336–6349 (2021).

19. Fonken, L.K., et al. Diminished circadian rhythms in hippocampal microglia may contribute to age-related neuroinflammatory sensitization. Neurobiol Aging 47, 102–112 (2016).

20. Cermakian, N., Westfall, S. & Kiessling, S. Circadian clocks and inflammation: reciprocal regulation and shared mediators. Arch Immunol Ther Exp (Warsz) 62, 303–318 (2014).

21. Nakazato, R., et al. The intrinsic microglial clock system regulates interleukin-6 expression. Glia 65, 198–208 (2017).

22. Marpegan, L., et al. Diurnal variation in endotoxin-induced mortality in mice: correlation with proinflammatory factors. Chronobiol Int 26, 1430–1442 (2009).

23. Sato, S., et al. A circadian clock gene, Rev-erbalpha, modulates the inflammatory function of macrophages through the negative regulation of Ccl2 expression. J Immunol 192, 407–417 (2014).

24. Imeri, L. & Opp, M.R. How (and why) the immune system makes us sleep. Nat Rev Neurosci 10, 199–210 (2009).

25. Leone, M.J., Marpegan, L., Duhart, J.M. & Golombek, D.A. Role of proinflammatory cytokines on lipopolysaccharide-induced phase shifts in locomotor activity circadian rhythm. Chronobiol Int 29, 715–723 (2012).

26. Group, T.H.s.D.C.R. A novel gene containing a trinucleotide repeat that is expanded and unstable on Huntington’s disease chromosomes. Cell 26;72(6), 971–983 (1993).

27. Politis, M., et al. Increased central microglial activation associated with peripheral cytokine levels in premanifest Huntington’s disease gene carriers. Neurobiol Dis 83, 115–121 (2015).

28. Morton, A.J., et al. Disintegration of the sleep-wake cycle and circadian timing in Huntington’s disease. J Neurosci 25, 157–163 (2005).

29. Loh, D.H., Kudo, T., Truong, D., Wu, Y.F. & Colwell, C.S. The Q175 Mouse Model of Huntington’s Disease Shows Gene Dosage- and Age-Related Decline in Circadian Rhythms of Activity and Sleep. Plos One 8 (2013).

30. Kudo, T., et al. Dysfunctions in circadian behavior and physiology in mouse models of Huntington’s disease. Exp Neurol 228, 80–90 (2011).

31. Aylward, E.H., et al. Onset and rate of striatal atrophy in preclinical Huntington disease. Neurology 63, 66–72 (2004).

32. Goodman, A.O., Morton, A.J. & Barker, R.A. Identifying sleep disturbances in Huntington’s disease using a simple disease-focused questionnaire. PLoS Curr 2, RRN1189 (2010).

33. Wood, T.E., et al. Mutant huntingtin reduction in astrocytes slows disease progression in the BACHD conditional Huntington’s disease mouse model. Hum Mol Genet 28, 487–500 (2019).

34. Savage, J.C., et al. Microglial physiological properties and interactions with synapses are altered at presymptomatic stages in a mouse model of Huntington’s disease pathology. J Neuroinflammation 17, 98 (2020).

35. Crotti, A., et al. Mutant Huntingtin promotes autonomous microglia activation via myeloid lineage-determining factors. Nat Neurosci 17, 513–521 (2014).

36. Ellrichmann, G., Reick, C., Saft, C. & Linker, R.A. The role of the immune system in Huntington’s disease. Clin Dev Immunol 2013, 541259 (2013).

37. Ni, J., et al. An impaired intrinsic microglial clock system induces neuroinflammatory alterations in the early stage of amyloid precursor protein knock-in mouse brain. J Neuroinflammation 16, 173 (2019).

38. Masuda, T., Sankowski, R., Staszewski, O. & Prinz, M. Microglia Heterogeneity in the Single-Cell Era. Cell Rep 30, 1271–1281 (2020).

39. Keren-Shaul, H., et al. A Unique Microglia Type Associated with Restricting Development of Alzheimer’s Disease. Cell 169, 1276–1290 e1217 (2017).

40. Dong, X.Y. & Cong, S.Y. Identification of differentially expressed genes and regulatory relationships in Huntington’s disease by bioinformatics analysis. Mol Med Rep 17, 4317–4326 (2018).

41. Vonsattel, J.P., Keller, C. & Cortes Ramirez, E.P. Huntington’s disease - neuropathology. Handb Clin Neurol 100, 83–100 (2011).

42. Harding, R.J. & Tong, Y.F. Proteostasis in Huntington’s disease: disease mechanisms and therapeutic opportunities. Acta Pharmacol Sin 39, 754–769 (2018).

43. Duennwald, M.L. & Lindquist, S. Impaired ERAD and ER stress are early and specific events in polyglutamine toxicity. Genes Dev 22, 3308–3319 (2008).

44. Browne, S.E., Ferrante, R.J. & Beal, M.F. Oxidative stress in Huntington’s disease. Brain Pathol 9, 147–163 (1999).

45. Relógio, A., et al. Tuning the mammalian circadian clock: robust synergy of two loops. PLoS Comput Biol 7, e1002309 (2011).

46. Curtis, A.M., Bellet, M.M., Sassone-Corsi, P. & O’Neill, L.A. Circadian clock proteins and immunity. Immunity 40, 178–186 (2014).

47. Conductier, G., Blondeau, N., Guyon, A., Nahon, J.L. & Rovere, C. The role of monocyte chemoattractant protein MCP1/CCL2 in neuroinflammatory diseases. J Neuroimmunol 224, 93–100 (2010).

48. Bjorkqvist, M., et al. A novel pathogenic pathway of immune activation detectable before clinical onset in Huntington’s disease. J Exp Med 205, 1869–1877 (2008).

49. Silver, A.C., Arjona, A., Walker, W.E. & Fikrig, E. The circadian clock controls toll-like receptor 9-mediated innate and adaptive immunity. Immunity 36, 251–261 (2012).

50. Lang, V., Ferencik, S., Ananthasubramaniam, B., Kramer, A. & Maier, B. Susceptibility rhythm to bacterial endotoxin in myeloid clock-knockout mice. Elife 10 (2021).

51. Fonken, L.K., et al. Stress-induced neuroinflammatory priming is time of day dependent. Psychoneuroendocrino 66, 82–90 (2016).

52. Hammond, T.R., et al. Single-Cell RNA Sequencing of Microglia throughout the Mouse Lifespan and in the Injured Brain Reveals Complex Cell-State Changes. Immunity 50, 253–271 e256 (2019).

53. Palpagama, T.H., Waldvogel, H.J., Faull, R.L.M. & Kwakowsky, A. The Role of Microglia and Astrocytes in Huntington’s Disease. Front Mol Neurosci 12, 258 (2019).

54. Wilton, D.K. & Stevens, B. The contribution of glial cells to Huntington’s disease pathogenesis. Neurobiol Dis 143, 104963 (2020).

55. Shin, J.Y., et al. Expression of mutant huntingtin in glial cells contributes to neuronal excitotoxicity. J Cell Biol 171, 1001–1012 (2005).

56. Petkau, T.L., et al. Mutant huntingtin expression in microglia is neither required nor sufficient to cause the Huntington’s disease-like phenotype in BACHD mice. Hum Mol Genet 28, 1661–1670 (2019).

57. Ben Haim, L., Carrillo-de Sauvage, M.A., Ceyzeriat, K. & Escartin, C. Elusive roles for reactive astrocytes in neurodegenerative diseases. Front Cell Neurosci 9, 278 (2015).

58. Crotti, A. & Glass, C.K. The choreography of neuroinflammation in Huntington’s disease. Trends Immunol 36, 364–373 (2015).

59. Ransohoff, R.M. How neuroinflammation contributes to neurodegeneration. Science 353, 777–783 (2016).

60. Wang, N., et al. Mapping brain gene coexpression in daytime transcriptomes unveils diurnal molecular networks and deciphers perturbation gene signatures. Neuron 110, 3318–3338 e3319 (2022).

61. Andrade-Navarro, M.A., et al. RNA Sequencing of Human Peripheral Blood Cells Indicates Upregulation of Immune-Related Genes in Huntington’s Disease. Front Neurol 11, 573560 (2020).

62. Mangiarini, L., et al. Exon 1 of the HD gene with an expanded CAG repeat is sufficient to cause a progressive neurological phenotype in transgenic mice. Cell 87, 493–506 (1996).

63. Jaitin, D.A., et al. Massively parallel single-cell RNA-seq for marker-free decomposition of tissues into cell types. Science 343, 776–779 (2014).

64. Keren-Shaul, H., et al. MARS-seq2.0: an experimental and analytical pipeline for indexed sorting combined with single-cell RNA sequencing. Nat Protoc 14, 1841–1862 (2019).

65. Thaben, P.F. & Westermark, P.O. Detecting rhythms in time series with RAIN. J Biol Rhythms 29, 391–400 (2014).

66. Carlucci, M., et al. DiscoRhythm: an easy-to-use web application and R package for discovering rhythmicity. Bioinformatics 36, 1952–1954 (2019).

67. Thaben, P.F. & Westermark, P.O. Differential rhythmicity: detecting altered rhythmicity in biological data. Bioinformatics 32, 2800–2808 (2016).

68. Huang, D.W., et al. The DAVID Gene Functional Classification Tool: a novel biological module-centric algorithm to functionally analyze large gene lists. Genome Biol 8, R183 (2007).

