## Supplemental material for "Single-cell variations of circadian clock and immune gene expression in microglia and neurodegeneration"

### Supplementary Information document

#### Supplementary Figure 1

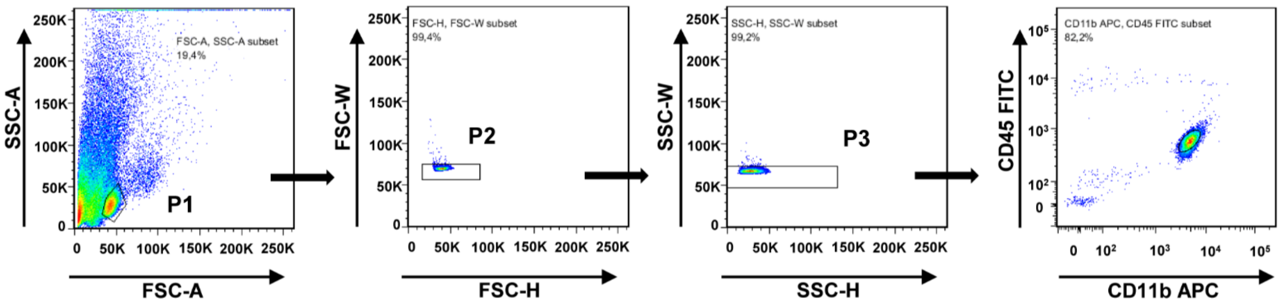


**Figure S1. Representative gating strategy for sorting microglia from 6- and 15-week-old R6/2 and WT mice by flow cytometry**

Cell population P1 was first separated from debris via FFC-A/SSC-A. Single cell gating was further performed with FSC-H/FSC-W to obtain population cells P2 after excluding extra wide or extra tall events that represent doublets or aggregated cells. Another single cell gating with SSC-H/SSC-W resulted in P3. Microglia were then sorted ($CD11b^{+}/CD45^{low}$), distinguishable by their CD45 expression level from macrophages ($CD11b^{+}/CD45^{high}$).

#### Supplementary Tables 1–8, Video 1

**Table S1**: Immune-relevant genes with differential rhythmicity identified between genotypes in 6-week-old R6/2 and control groups

**Table S2**: Immune-relevant genes with differential rhythmicity identified between genotypes in 15-week-old R6/2 and control groups

**Table S3**: KEGG data-based immune pathways related to mouse module

**Table S4**: List of genes obtained after intersection between the differentially rhythmic genes identified in 6-week-old R6/2 and control groups and genes involved in the murine KEGG data-based immune pathways

**Table S5**: List of genes obtained after intersection between the differentially rhythmic genes identified in 15-week-old R6/2 and control groups and genes involved in the murine KEGG data-based immune pathways

**Video1**: 3D-Reconstruction of microglia using Imaris software

**Table S6**: Rhythmicity analysis results of RNAscope candidates

**Table S7**: Differential rhythmicity results of RNAscope candidates (R6/2 versus WT mice)

**Table S8**: Differential rhythmicity results of RNAscope candidates (between different tissues of same genotype)
